# DARPins recognizing mTFP1 as novel reagents for *in vitro* and *in vivo* protein manipulations

**DOI:** 10.1101/354134

**Authors:** M. Alessandra Vigano, Dimitri Bieli, Jonas V. Schaefer, Roman Peter Jakob, Shinya Matsuda, Timm Maier, Andreas Plückthun, Markus Affolter

## Abstract

Over the last few years, protein-based affinity reagents have proven very helpful in cell and developmental biology. While many of these versatile small proteins can be expressed both in the intracellular and extracellular milieu in cultured cells and in living organisms, they can also be functionalized by fusing them to different protein domains in order to regulate or modulate their target proteins in diverse manners. For example, protein binders have been employed to degrade, trap, localize or enzymatically modify specific target proteins. Whereas binders to many endogenous proteins or small protein tags have been generated, also several affinity reagents against fluorescent proteins have been created and used to manipulate target proteins tagged with the corresponding fluorescent protein. Both of these approaches have resulted in improved methods for cell biological and developmental studies. While binders against GFP and mCherry have been previously isolated and validated, we now report the generation and utilization of designed ankyrin repeat proteins (DARPins) against the monomeric teal fluorescent protein 1 (mTFP1). Here we use the generated DARPins to delocalize Rab proteins to the nuclear compartment, in which they cannot fulfill their regular functions anymore. In the future, such manipulations might enable the production of acute loss-of-function phenotypes in different cell types or living organisms based on direct protein manipulation rather than on genetic loss-of-function analyses.

**Summary statement:** Structural characterization of two novel DARPins (designed ankyrin repeat proteins) recognizing the monomeric teal fluorescent protein 1 (mTFP1) and their functionalization for protein manipulation strategies in cultured cells and potentially in living organisms.

## Introduction

Over the last decades, much has been learned about the role of different proteins in controlling cell proliferation, cell movement, cell determination and cell differentiation, both in cell culture and during the development of multicellular animals. To a large extent, such knowledge was gained by comparing the behaviour of wild-type and mutant individuals, starting with large-scale genetic screens (Brenner, 1974; Nüsslein-Volhard and Wieschaus, 1980). Later, these approaches were complemented with reverse genetic approaches, which allowed for loss- and gain-of-function studies that could be controlled with regard to developmental time and to the targeted tissue (Anderson et al., 2017; Housden et al., 2017; Nagarkar-Jaiswal et al., 2015; Nagarkar-Jaiswal et al., 2017; Venken et al., 2011; Yamamoto et al., 2014). To further increase the possibilities to study protein function, RNAi- and morpholino-oligonucleotide-based methods were used to induce a reduction in protein levels. While off-target effects have to be taken into account when studying protein function with these methods, these approaches, in particular RNAi, allow for time-and tissue-controlled, genome-wide loss-of-function analyses (Housden et al., 2017). In all of these approaches, the level of the target protein is reduced either by the lack of function of the gene, or by decreased levels of mRNA. However, when studying proteins with a particularly long half-life, or in cases of maternal contribution of proteins in early embryos, it might be very difficult or even impossible to deplete the protein of interest and analyse its contribution to cellular or organismal function.

To circumvent this problem, different approaches allowing the direct manipulation of protein levels were recently developed. A number of different methods were established to degrade proteins in an inducible fashion (Banaszynski et al., 2006; Bonger et al., 2011; Chung et al., 2015; Harder et al., 2008; Natsume et al., 2016), or to remove proteins from their place of action and thereby inactivating or preventing their functions in their native environment, (anchor away (Haruki et al., 2008) and knocksideways (Robinson et al., 2010)). In the last few years, optogenetic tools were designed to regulate protein activity or protein dimerization; these light-controllable tools are now used in cell and developmental biology to regulate or manipulate protein function in a more controllable fashion (Buckley et al., 2016; Guglielmi et al., 2016; Rost et al., 2017).

Recently, another, somewhat different approach has emerged, which allows targeting and manipulating proteins in several ways and in a more systematic manner. Using small protein scaffolds, it became possible to screen for and isolate binders against proteins of interest, post-translational modifications of proteins or against protein tags such as fluorescent proteins (Beghein and Gettemans, 2017; Bieli et al., 2016; Harmansa and Affolter, 2018; Helma et al., 2015; Plückthun, 2015; Sha et al., 2017). Such protein binders have been used extensively as crystallization chaperones in structural biology (Batyuk et al., 2016; Manglik et al., 2017), as high-affinity reagents or sensors (Borg et al., 2015; Braun et al., 2016; Kummer et al., 2013; Rothbauer et al., 2008; Trinkle-Mulcahy et al., 2008), as detection reagents in light and super resolution microscopy (Pleiner et al., 2015; Ries et al., 2012) and for targeting medically relevant intracellular proteins (Boldicke, 2017; Grebien et al., 2011; Koide et al., 2012).

Furthermore, functionalized protein binders have emerged as versatile tools to target and manipulate proteins *in vivo* for developmental studies. In most of these studies, binders against fluorescent proteins were used to target proteins of interest fused to the corresponding fluorescent proteins. Such functionalized binders allowed the visualization, the degradation, the delocalization or the chemical modification of the specific target, and thereby provide insight into the functional roles of proteins in developmental processes (reviewed in (Beghein and Gettemans, 2017; Helma et al., 2015; Plückthun, 2015; Sha et al., 2017)).

In cell and developmental biology, it is now a standard procedure to use several fluorescent proteins simultaneously to analyse complex processes *in vivo* and in real time. It would therefore be valuable to have specific binders against many different fluorescent proteins in order to be able to manipulate and/or follow different proteins simultaneously. At present, only a limited number of binders for GFP (green fluorescent protein) and mCherry have been isolated and characterized (Brauchle et al., 2014; Fridy et al., 2014; Kubala et al., 2010; Moutel et al., 2016).

Here, we report the selection of designed ankyrin repeat proteins (DARPins) (Plückthun, 2015) recognizing mTFP1 (monomeric teal fluorescent protein 1). We characterized these binders both biochemically and biophysically and determined the three-dimensional structure of one DARPin-mTFP1 complex. *In vivo* functionality of anti-mTFP1 DARPins was demonstrated in delocalization experiments using Rab proteins. In the future, such manipulations could enable the generation of acute loss-of-function phenotypes in different cell types based on protein manipulation rather than genetic loss-of-function analyses.

## Results

We have previously reported the isolation and characterization of DARPins recognizing GFP and mCherry, including “clamp” constructs (Brauchle et al., 2014; Hansen et al., 2017). To further increase the number of orthogonal reagents available to selectively target fluorescent fusion proteins, we wanted to generate DARPins against a fluorescent protein absorbing and emitting light in a different range of the light spectrum. We decided to target mTFP1 since at the time it represented the brightest monomeric protein in the blue-green spectrum (Ai et al., 2006; Ai et al., 2008). mTFP1 was produced recombinantly in a prokaryotic expression system and used to select DARPins against this target.

### Selection and *in vitro* characterization of mTFP1-binding DARPins

To generate suitable DARPin binders, streptavidin-binding peptide (SBP)-tagged mTFP1 was immobilized on streptavidin beads and used as a target for DARPin selections by employing multiple rounds of Ribosome Display (Dreier and Plückthun, 2012; Plückthun, 2012). In each round, the target concentration presented on magnetic streptavidin beads was decreased while the washing stringency was simultaneously increased to enrich for binders with high affinities. After four rounds of selection, the enriched pool was cloned into an *E. coli* expression vector, allowing the production of both N-terminally His8- and C-terminally FLAG-tagged DARPins. Nearly four hundred colonies of transformed *E. coli* were picked and the encoded DARPins were expressed in small scale. Bacterial crude extracts were subsequently used in ELISA screenings, detecting the binding of candidate DARPins to streptavidin-immobilized mTFP1 by employing a FLAG-tag based detection system (data not shown). The top 30 candidates from this initial ELISA screens were analysed in more detail, testing their binding to streptavidin-immobilized mTFP1 and comparing them to the interaction with streptavidin alone. Of these analysed clones, only two candidates, named 1238_E11 and 1238_G01, showed a specific binding to mTFP1 while the majority of previous hits seemed to also be interacting with free streptavidin. This very unusually low number, in this experiment, compared to the usual 50-200 specific binders, is almost certainly a consequence of attempting to immobilize the target via a streptavidin-binding peptide, instead of the usual biotin (see Discussion). The sequence of the two selected DARPins is shown in Supplementary Figure S1A.

### Specificity, Affinity and Epitopes of Top-two DARPin candidates

To test whether the selected DARPins 1238_E11 and 1238_G01 are specific for mTFP1, titration ELISAs against mTFP1, two other fluorophores (GFP and mCherry) and the Maltose Binding Protein (MBP) as an unrelated protein target were performed. As shown in Figure 1A, both DARPins clearly displayed a specificity for mTFP1, with apparent affinities in the low nM range (Figure 1A, blue curves). However, it was also noted that DARPin 1238_E11 (left panel in Figure 1A) had both a higher apparent affinity to the target and a higher background binding compared to DARPins 1238_G01 (right panel in Figure 1A).

**Figure 1.**
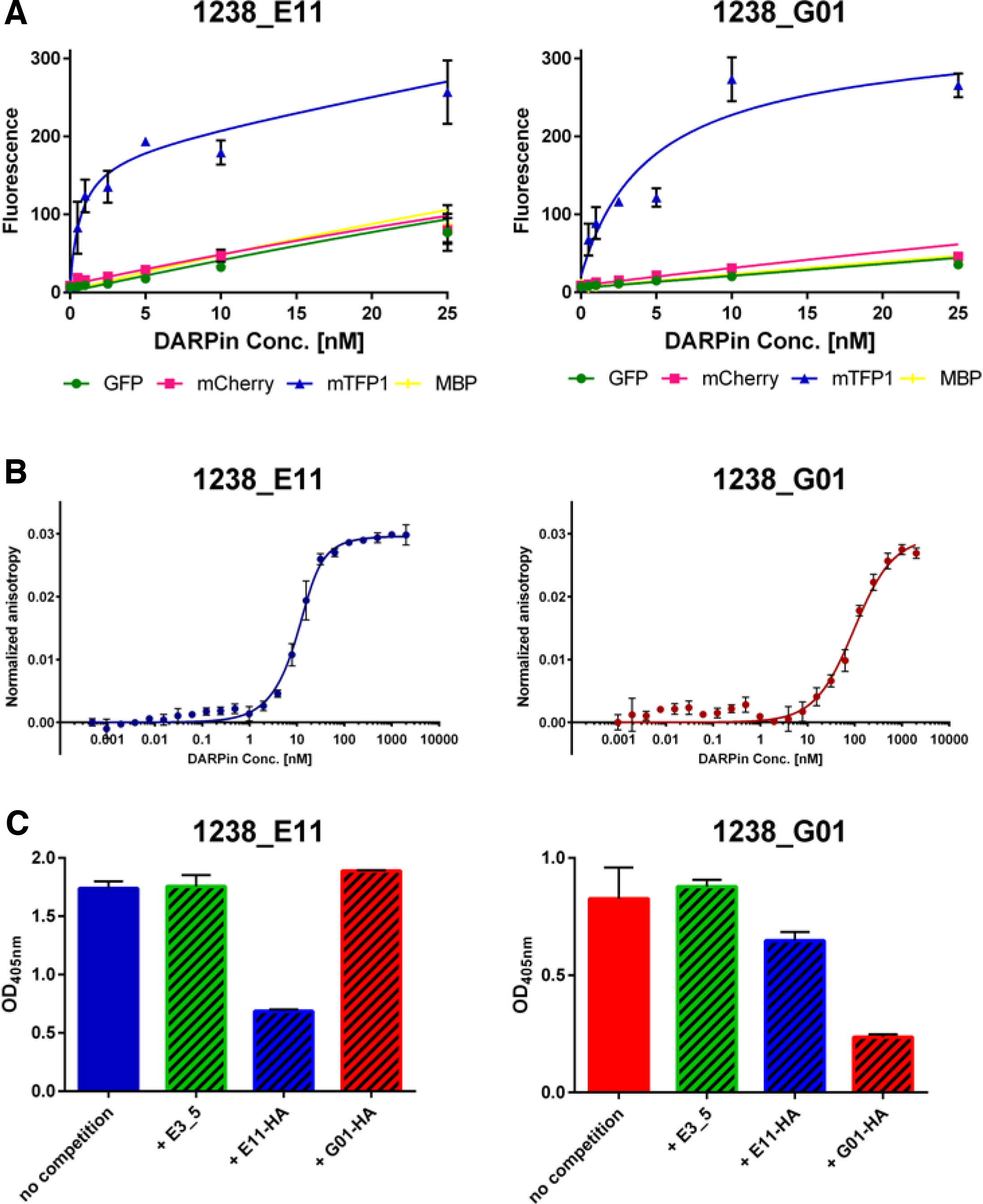
Biochemical analyses of selected anti-mTFP1 DARPins. **(A)** Titration ELISA: DARPins 1238_E11 (left panel) and 1238_G01 (right panel) show specific binding to mTFP1 over control surfaces (GFP, mCherry and MBP). **(B)** Fluorescence anisotropy measurements of 1238_E11 (left panel) and 1238_G01 (right panel) reveal high affinities with K_D_ values of 3 nM and 88 nM, respectively. **(C)** Epitope blocking ELISA. Immobilized mTFP1 was incubated with a mixture of HA-and FLAG-tagged DARPins with a relative 1:5 ratio (100 nM of the FLAG-tagged DARPins and 500 nM of the HA-tagged competitor) to analyse the influence of the competitor on the original signal (shown on left side, named “no competition“).

The affinities of these two DARPins were then measured by fluorescence anisotropy as previously described (Brauchle et al., 2014) (Figure 1B). For DARPins 1238_E11 (left panel), a high affinity with a K_D_ value of 3 nM was determined, while the affinity of 1238_G01 was found to be lower with a value of about 88 nM.

To analyse whether the two DARPins recognize different, non-overlapping epitopes on mTFP1, competition/epitope blocking ELISAs were performed. Streptavidin-immobilized mTFP1 was incubated with a mixture of FLAG- and HA-tagged DARPins, with the HA-tagged binders being present at a five-fold higher concentration over the finally detected FLAG-tagged binders. As shown in the left panel in Figure 1C for DARPin 1238_E11, the presence of neither the non-binding DARPin E3_5 (green striped bar), nor of the HA-tagged DARPin 1238_G01 (red striped bar) had reduced the FLAG-based detected signals. However, when the identical DARPin binder was present as an HA-tagged variant (blue striped bar), the detected signal was clearly diminished. Similar results were obtained for DARPin 1238_G01 as shown in the right panel of Figure 1C. These data suggest that the two anti-mTFP1 DARPins bind to different epitopes on their target.

### Structural Analysis of the mTFP1/DARPin 1238_E11 Complex

In order to understand the basis for specificity and structural changes in the binding of the selected DARPins to mTFP1, the crystal structures of the mTFP1/DARPin 1238_E11 complex were determined (Figure 2, Supplementary Figure S1). The complex with DARPins 1238_G01 did not crystallize, but the structure of both isolated DARPins was determined as well (1238_G01, 1.6 Å resolution, R/R_free_ of 16/18) and (1238_E11, 2.1 Å resolution, R/R_free_ of 17/21).

**Figure 2.**
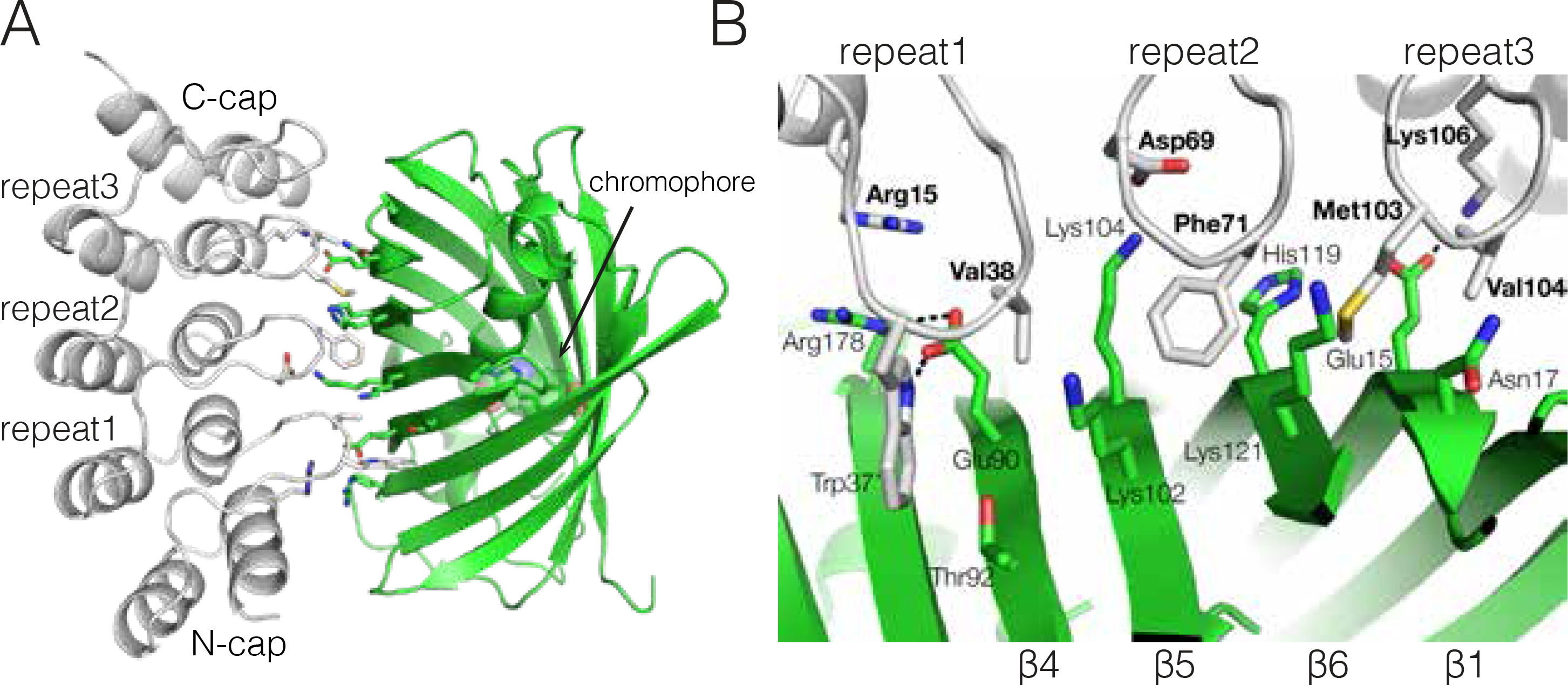
Structure of the mTFP1/mTFP1-DARPin 1238_E11 complex. **(A)** Cartoon representation of mTFP1 (green) and DARPin 1238_E11 (grey). The mTFP1 chromophore is shown in sphere representation. **(B)** Close-up view of the binding interface of mTFP1 (green) and DARPin 1238_E11 (grey). DARPin residues are labelled in bold; hydrogen bonding interactions are indicated by black dashed lines.

The mTFP1/DARPin 1238_E11 complex was obtained by mixing mTFP1 with a 1.2-fold excess of DARPin 1238_E11 and subjecting the mixture to gel filtration. Gel filtration analysis indicated a homogeneous 1:1 complex, which was used for crystallization as described in Materials and Methods. We determined the crystal structure of the mTFP1/-DARPin 1238_E11 complex in two space groups, to 1.6 Å (P6_5_22) and 1.85 Å (C2) resolution, respectively (Figure 2A). Both complex structures are structurally identical (C_α_ r.m.s.d. of 0.3 Å): The overall fold of mTFP1 exhibits the typical β-can motif of the GFP family (Yang et al., 1996) and undergoes only minor structural changes upon DARPin binding (Ai et al., 2006; Henderson et al., 2007). Similarly, upon mTFP1 binding, DARPin 1238_E11 remains structurally unchanged and exhibits only minor differences in the N-Cap region (Supplementary Figure S1C; C_α_ r.m.s.d. of 0.45 Å). DARPin 1238_E11 binds into a cavity along mTFP1 including β-strands 1, 4, 5 and 6, comprising a protein-protein binding surface of 790 Å^2^ (Krissinel and Henrick, 2007) and thus similar in size compared to other known DARPin protein complexes (Gilbreth and Koide, 2012; Sennhauser and Grütter, 2008). Residues from the N-Cap and the three DARPin repeats make specific contacts with mTFP1 (Figure 2B). Interactions involve hydrophobic DARPin residues Trp37, Val38, Phe71, Met103 and Val104, two salt bridges (Asp69-Lys104 and Lys106-Glu15) and an arginine-arginine paring (Magalhaes et al., 1994; Zhang et al., 2013)(Arg15-Arg178) stabilized by Glu90 (Figure 2B).

### anti-mTFP1 DARPins bind mTFP1 fusion proteins in cultured cells

Since many developmental and cellular processes take place in intracellular compartments, we tested the expression and functionality of the anti-mTFP1 DARPins within cells. In order to visualize the two DARPins, we fused 1238_E11 and 1238_G01 to different additional fluorescent proteins (mCherry or YPet) and used transient transfection in HeLa cells as a model system to test their properties (Brauchle et al., 2014; Moutel et al., 2016). The two DARPins 1238_E11 and 1238_G01 were transfected either alone or in combination with mTFP1 fusion constructs which localize to specific compartments or specific membranes of the cells. When no mTFP1 was co-expressed, the DARPin-mCherry fusions alone were uniformly expressed and localized in both the cytoplasm and nucleus of transfected cells, with a stronger accumulation in the nuclei (Figure 3 panels of A and B). In contrast, a mitochondrial mTFP1 “bait” alone (see Materials and Methods for a description of the mito-mTFP1 construct) was visible at the mitochondria of the transfected cells (Figure 3, panels of C), due to the fusion with the N-terminal domain of the outer mitochondrial membrane protein MitoNEET (CISD1), which is N-terminally anchored to the outer membrane and exposed to the cytoplasmic outside of the mitochondria (Colca et al., 2004; Wang et al., 2017), (see Supplementary Figure S4 for a schematic representation of the fusion proteins and Supplementary file S1 for their amino acid sequences). When these two constructs coding for mito-mTFP1 and DARPin-mCherry and were co-transfected, we observed a clear mitochondrial co-localization (Figure 3, panels of D and E), demonstrating binding and recruitment of DARPin-mCherry to the mitochondrial surface via the mitochondrial mTFP1-bait.

**Figure 3.**
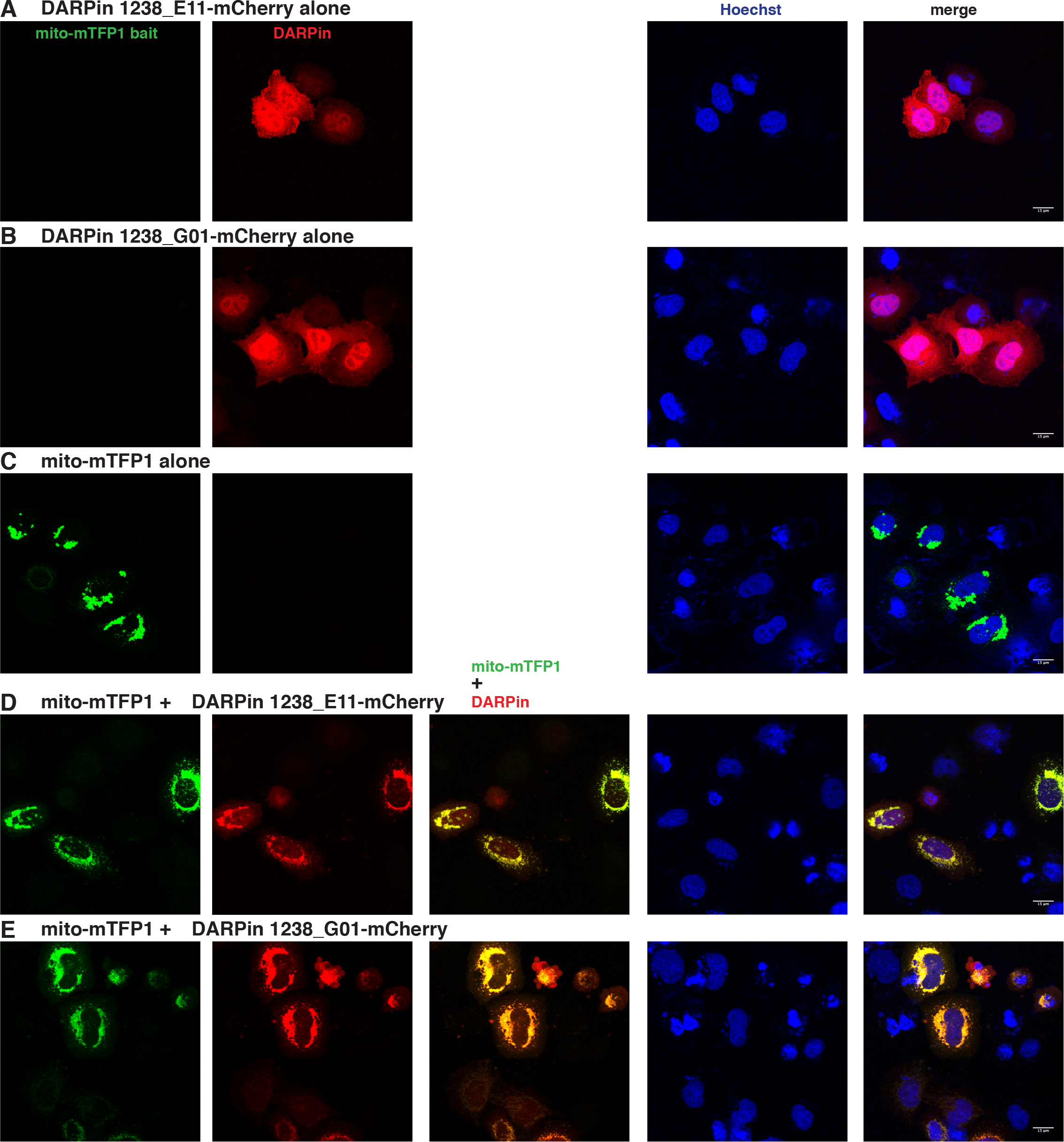
Intracellular binding of anti-mTFP1-DARPins. Confocal images of HeLa cells transiently transfected **(A)** with pCMV-DARPin 1238_E11-mCherry alone, **(B)** pCMV-DARPin 1238_G01-mCherry alone, **(C)** pMITO-mTFP1 alone, **(D)** the combination of pMITO-mTFP1 and pCMV-DARPin 1238_E11-mCherry**, (E)** pMITO-mTFP1 and pCMV-DARPin 1238_G01-mCherry. The first column represents the mTFP1 channel (green), the second column is the mCherry channel (red), the third column is the overlay of the two channels, showing the mitochondrial colocalization (indicated in yellow) of the mito-mTFP1 bait with the respective DARPin, the fourth column represents the nuclear Hoechst staining (blue) and the fifth column is the merge of all three channels (with the scale bar in white (15 μm) on the bottom right corner). Images were taken 24 hours post transfection. Transfected constructs are indicated on top of each row and the different channels are indicated inside the panels of the first row. The figures are from a representative experiment, performed at least three times.

Nonetheless, it has to be noted that not all the DARPin molecules were recruited to the mitochondria, as seen by residual mCherry signal in the cytoplasm, presumably because of the limited number of CISD1 binding partners at the mitochondrial surface. Also, varying the ratio of the transfected DNAs did not change the amount of DARPins observed at the mitochondria (data not shown). Furthermore, the different binding affinities of the two DARPins are also not reflected in this type of experiment: it appears that these affinities are sufficient to recruit both DARPins to mitochondria in a very similar way.

To ensure that the specific binding properties of the anti-mTFP1 DARPins would work in many contexts, we also tested them for co-localization with mTFP1-baits expressed in different cellular sites (Supplementary Figure S2) Thus, we re-localized the DARPin-mCherry proteins to the nuclear compartment via binding to a histone H2B (H2B)-mTFP1 fusion (Supplementary Figure S2, panels of first row). Indeed, a strong nuclear co-localization with this bait was observed for both DARPins (Supplementary Figure S2, panels of second and third row).

Finally, we tested a membrane-localizing mTFP1 bait: mTFP1-CAAX (mTFP1 with the C-terminal HRas farnesylation motif CVLS) together with the two anti-mTFP1 DARPins. Also in this combination, we observed a clear co-localization of the mCherry signal from both DARPin-mCherry fusion proteins with the mTFP1 signal at the cell membrane (Supplementary Figure S2, panels of fourth, fifth and sixth row).

To confirm that the mitochondrial and nuclear co-localizations with the respective mTFP1 baits were truly mediated by direct binding of the DARPins to mTFP1 and not by an unspecific mTFP1-mCherry interaction, we repeated the co-localization experiments in HeLa cells, fusing the DARPins to YPet, another fluorescent protein (Nguyen and Daugherty, 2005). The results shown in Supplementary Figure S3 confirmed the specific binding of the DARPins-YPet to the mTFP1 baits both in the nuclei and at the mitochondria.

### anti-mTFP1 DARPins can be functionalized for *in vivo* relocalization experiments

As a next step, we tested whether the anti-mTFP1 DARPins could bind and thereby re-localized a mTFP1-fusion protein when stably tethered to a specific subcellular compartment. For this purpose, we created a mTFP1-Rab5c fusion construct, where Rab5c would be expected to be mainly localized to the cytoplasmic face of the early endosome via its lipid anchors. In an experimental setup similar to the one previously used for an anti-GFP-DARPin (Brauchle et al., 2014), we added a CAAX motif to both DARPin-YPet fusion proteins in order to tether them to the cell membrane, facing towards the cytoplasm. As anticipated, these DARPin-YPet-CAAX fusions were localized mostly along the plasma membrane of the transfected cells (Figure 4, panels of B and C). When transfected alone, the mTFP1-Rab5c fusion construct showed both a diffuse and a “vesicle-like” distribution, especially in the perinuclear region, but it was never observed inside the nucleus (Figure 4 and Figure 5, panels of A). In contrast, upon cotransfection with either of the membrane-tethered DARPins, mTFP1-Rab5c was re-localized to the plasma membrane (Figure 4, panels of A vs. D and E). Although some mTFP1-Rab5c was still visible in the perinuclear region, most of it was re-localized at the plasma membrane under these cotransfection conditions, thus demonstrating that both anti-mTFP1 DARPins can be functionalized for in vivo re-localization experiments.

**Figure 4.**
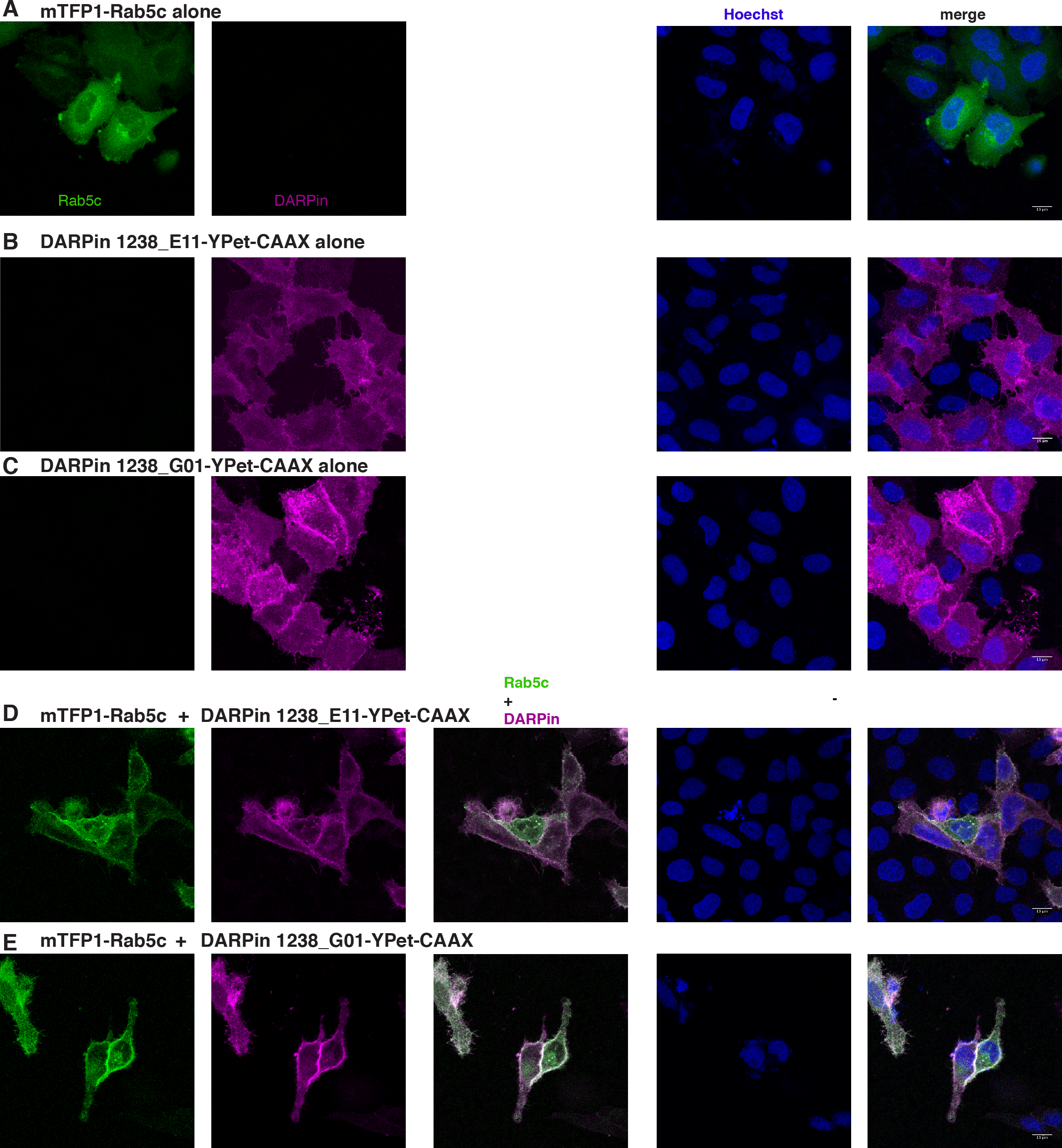
Rab5c recruitment to plasma membrane with anti-mTFP1-DARPins. Confocal images of HeLa cells transiently transfected **(A)** with pcDNA3-mTFP1-Rab5c alone, **(B)** pCMV-DARPin 1238_E11-YPet-CAAX alone**, (C)** pCMV-DARPin 1238_G01-YPet-CAAX alone, **(D)** the combination of pcDNA3-mTFP1-Rab5c and pCMV-DARPin 1238_E11-YPet-CAAX, **(E)** pcDNA3-mTFP1-Rab5c and pCMV-DARPin 1238_G01-YPet-CAAX. The first column represents the mTFP1 channel (green), the second column is the YPet channel (magenta), the third column is the overlay of the two channels, showing the recruitment of mTFP1-Rab5c to the plasma membranes, the fourth column represents the nuclear Hoechst staining (blue) and the fifth column is the merge of all three channels (with the scale bar in white (15 μm) on the bottom right corner). Images were taken 24 hours post transfection. Transfected constructs are indicated on top of each row and the different channels are indicated inside the panels of the first row. The figures are from a representative experiment, performed at least three times.

### Reverse anchor-away with protein binders

In the previous experiment, we demonstrated the use of DARPins to actively intervene into a cellular process by re-localizing Rab5c from its perinuclear localization to the plasma membrane. Another way to use the intracellular protein binders would be the redistribution of a protein to a place in the cell where it can no longer exert its native function (Haruki et al., 2008; Robinson et al., 2010). We used again mTFP1-tagged Rab5c together with the anti-mTFP1-DARPins, with the ultimate aim of removing Rab proteins from their natural subcellular site of action in the cytoplasm. Therefore, we fused the anti-mTFP1 DARPins-YPet to histone H2B to stably anchor them in the nucleus. Upon transfection of HeLa cells, the expression of the two DARPin-YPet-H2B fusions was exclusively nuclear, with a stronger accumulation in nucleoli or other unspecified nuclear bodies, as shown in Figure 5 (panels of B and C). Cotransfection of mTFP1-Rab5c and either of the DARPin-YPet-H2B fusions clearly brought most of the mTFP1-Rab5c signal into the nuclei of the cotransfected cells, indicating active removal of mTFP1-Rab5c from its site of action, the cytoplasmic/vesicular compartment (see panels of A vs. D and E in Figure 5).

**Figure 5.**
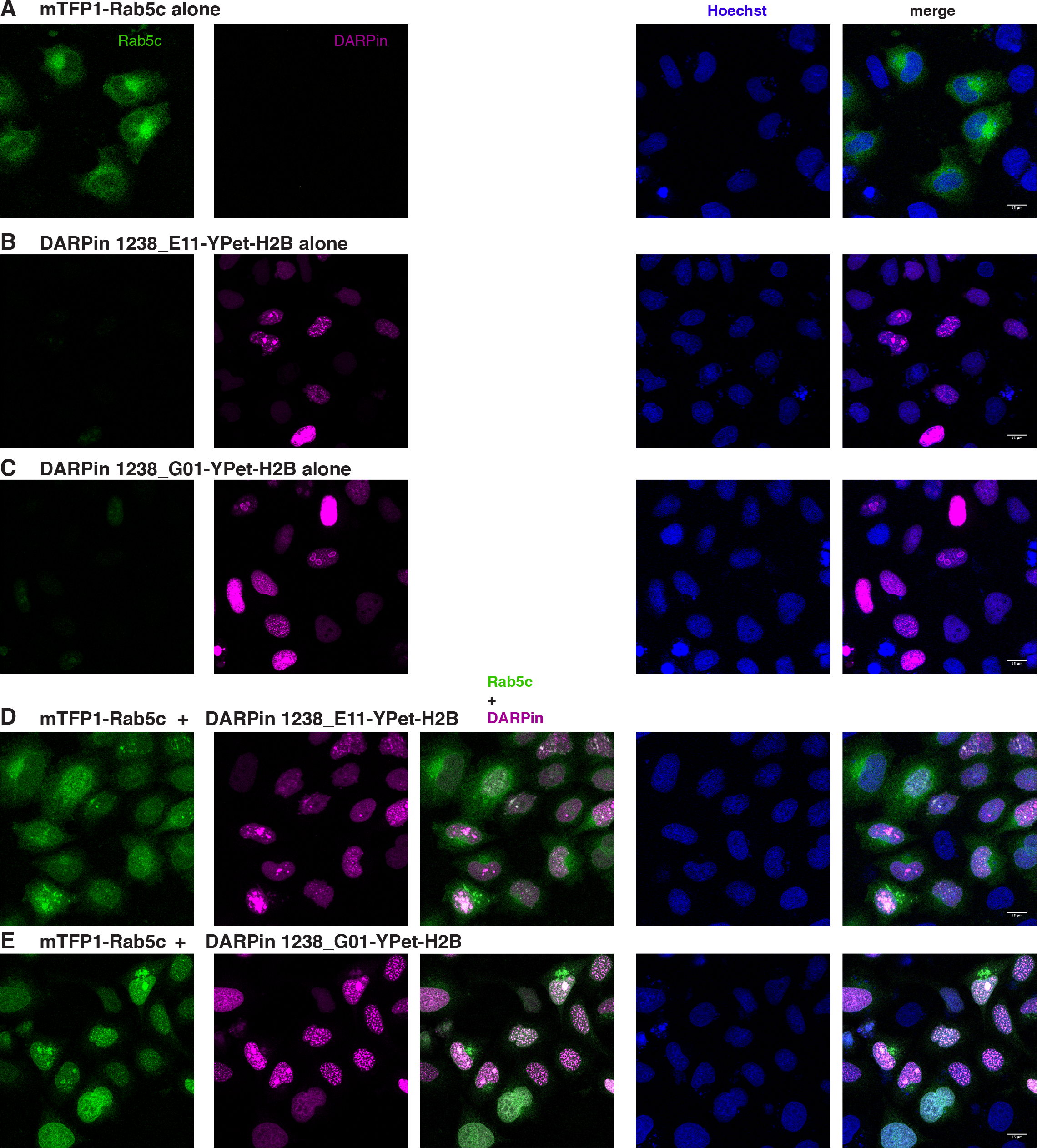
Rab5c trapping in the nucleus with anti-mTFP1-DARPins. Confocal images of HeLa cells transiently transfected with **(A)** pcDNA3-mTFP1-Rab5c alone, **(B)** pCMV-DARPin 1238_E11-YPet-H2B alone, **(C)** pCMV-DARPin 1238_G01-YPet-H2B alone, **(D)** the combination of pcDNA3-mTFP1-Rab5c and pCMV-DARPin 1238_E11-YPet-H2B, **(E)** pcDNA3-mTFP1-Rab5c and pCMV-DARPin 1238_G01-YPet-H2B. The first column represents the mTFP1 channel (green), the second column is the YPet channel (magenta), the third column is the overlay of the two channels, showing the recruitment of mTFP1-Rab5c to the nuclei, the fourth column represents the nuclear Hoechst staining (blue) and the fifth column is the merge of all three channels (with the scale bar in white (15 μm) on the bottom right corner). Images were taken 24 hours post transfection. Transfected constructs are indicated on top of each row and the different channels are indicated inside the panels of the first row. The figures are from a representative experiment, performed at least three times.

## Discussion

### Isolation and characterization of novel reagents against mTFP1

Protein binders against fluorescent proteins are valuable tools in biochemical research. Here, we report the selection of DARPins recognizing mTFP1, one of the brightest and most photostable FPs in the blue-green spectrum. While the two DARPins 1238_E11 and 1238_G01 proved to be of high value for the subsequent cellular assays, we were surprised to find a rather limited number of hits in our primary screening that satisfied our precondition of a signal of 40-fold over background. This number of hits is much lower than that in other selections performed in parallel, where typically 50 to 250 hits are found (data not shown) and this is most likely caused by the limited affinity of the streptavidin-binding peptide (SBP) to streptavidin (Keefe et al., 2001). Indeed, the immobilization strategy and washing steps in the performed Ribosome Display were previously optimized for the immobilization via biotinylated target proteins, and biotin binds essentially irreversibly to the streptavidin beads, while the SBP is washed away. Furthermore, the high percentage of identified background binders (i.e. DARPins binding to streptavidin and not mTFP1) is most likely caused by the fact that, with the immobilization via SBP, the routinely performed alternations between streptavidin and neutravidin between different rounds of Ribosome Display could not be employed, as SBP only binds to streptavidin, but not neutravidin. Therefore, we cannot recommend replacing biotinylation by streptavidin-binding peptides. Nevertheless, the two binders identified proved to be specific, having high affinity and binding to different, non-overlapping epitopes on mTFP1. This last property may render these two anti-mTFP1-DARPins suitable for a sandwich pair (Hansen et al., 2017).

### Structure of the mTFP1-DARPin complex

Using X-ray crystallography, we obtained a crystal structure for the tightest-binding mTFP1 binding DARPin 1238_E11 in complex with mTFP1. Comparison to the crystal structures of the individual proteins showed no significant structural changes for both proteins upon complex formation. DARPin 1238_E11 binds along the barrel of mTFP1, using hydrophobic and charge complementation (Figure 2). Anti-mTFP1 DARPin 1238_E11 and DARPin 1238_G01 differ in 13 out of 21 selected library residues (Supplementary Figure S1A), suggesting alternative binding epitopes, in line with the ELISA experiments (Figure 1C). Recently, the structures of the complexes of GFP with DARPin 3G61 (PDB ID: 5MAD; Supplementary Figure S1D) and DARPin 3G124nc (PDB ID: 5MA6; Supplementary Figure S1E) were determined (Hansen et al., 2017). GFP and mTFP1 show 27% identity and 1.4 Å C_α_ rmsd.

DARPin 3G61 and DARPin 3G124nc have an affinity to GFP in a similarly low nM range as DARPin 1238_E11 to mTFP1 (Brauchle et al., 2014), but use distinct, non-overlapping epitopes to bind to the barrel of the fluorescent protein (Supplementary Figure S1D and E). This comparison shows how the beta-barrel of the autofluorescent proteins can be bound from different sides by the DARPin paratopes.

### “*in vivo*” performance of the anti-mTFP1-DARPins

Next to other beneficial properties, DARPins fold very well and thus also display a high solubility in the intracellular milieu (Plückthun, 2015). Indeed, both of the selected anti-mTFP1-DARPins are well distributed inside the cell and do not form aggregates, even when overexpressed under the control of the CMV promoter and/or when fused to different fluorescent proteins (mCherry and YPet) (Figure 3 and Supplementary Figure S3). Furthermore, they are able to recognize and specifically bind mTFP1 *in vivo* in different subcellular compartments (Figure 3 and Supplementary Figure S2). Despite having different binding affinities, both DARPins, 1238_E11 and 1238_G01, bind in a very similar way inside the cell, suggesting that both affinities are sufficient. More importantly, both these binders can be “functionalized” in order to recruit an overexpressed mTFP1-Rab5c fusion protein to the plasma membrane (Figure 4) or to trap it into the nucleus (Figure 5), with likely different biological consequences. Therefore, these DARPins can now be employed in many different applications both *in vitro* and *in vivo*.

### Use of mTFP1 binders in cell and developmental biology

The number of different fluorescent proteins, which vary in their absorption and emission spectrum as well as in other properties (stability, photobleaching and their ability to be photoconverted or photoactivated), has steadily increased over the last decades (Rodriguez et al., 2017). In order to further manipulate the *in vivo* function of proteins of interests fused to such fluorescent moieties, it would help to have an equally diverse collection of small-protein binders recognizing these fluorescent proteins. The two DARPins we report here bind specifically and with high affinity to mTFP1. The use of these novel reagents will now allow performing more complex experiments, both in cultured cells and in multicellular organisms. Similar to the large number of different optogenetic tools developed over the past years (Rost et al., 2017), the availability of a battery of binders to different fluorescent proteins would allow for a multiparametric approach for imaging and manipulation. For example, we intend to use mTFP1-recognizing DARPins to mislocalize Rab proteins in the zebrafish vasculature and follow the behaviour of other cellular processes (e.g. trafficking of luminal and/or junctional proteins) using available marker proteins fused to GFP and red fluorescent proteins (mCherry or mKate2); for the latter two fluorescent proteins, there is a considerable number of lines available encoding fusion proteins that can be used. Ideally, the protein/gene of interest would be fused to mTFP1 at the endogenous locus in order to be expressed under the control of the endogenous regulatory sequences. The generation of such lines in zebrafish will take some time and therefore goes beyond the scope of this study.

In summary, the novel DARPins we characterize at the structural and functional level in this study contribute to the growing toolbox of protein-directed binder modules that can be used in a large variety of applications (Beghein and Gettemans, 2017; Helma et al., 2015; Plückthun, 2015) and thus should be of great value for the scientific community.

## Materials and Methods

### Protein expression and purification

DARPin protein constructs were cloned into pQiq vectors, overexpressed in *E. coli* XL1Blue cells and purified as described previously (Brauchle et al., 2014). For the expression of mTFP1, the gene fragment was cloned into the expression plasmid pRSFDuet-1 (Novagen), between BamHI and EcoRI sites, to generate N-terminally hexa-His-SBP-tagged mTFP1 fusion protein (Keefe et al., 2001). The amino acid sequence of these tagged proteins is provided in Supplementary file S1. The protein was overexpressed in *E. coli* BL21(DE3). After lysis of the cells in 50 mM Hepes/NaOH, 250 mM NaCl, pH 7.4, 40 mM imidazole with a sonicator and centrifugation, the proteins were purified by immobilized metal-affinity chromatography on a Ni-NTA column. For structural studies, purified mTFP1 and DARPin were mixed in a 1.2:1 ratio and then subjected to size-exclusion chromatography in 20 mM Hepes/NaOH pH 7.4 on a Superdex S75 column (GE Healthcare). The protein complex-containing fractions were pooled and concentrated in Amicon Ultra units (Millipore).

### ELISA and fluorescence anisotropy (FA) assay

Black 384-well Maxisorp plates (Nunc) were coated with 20 μl streptavidin (66 nM in PBS) overnight and blocked the next day with 100 μl PBS-TB (PBS containing 0.1% Tween-20 and 0.2% bovine serum albumin). After washing three times with 100 μl PBS-T, wells were coated with 20 μl of either *in vivo* biotinylated GFP, mCherry or MBP (maltose binding protein) or SBP-tagged mTFP1 at a concentration of 100 nM. Subsequently, 20 μl purified FLAG-tagged anti-mTFP1 DARPins 1238_E11 and 1238_G01 were applied in concentrations ranging from 1 to 50 nM and incubated for 1 h. Following another incubation with a horse radish peroxidase (HRP)-conjugated anti-FLAG antibody (Sigma-Aldrich #A8592, 1: 2,500) for 1 h, bound DARPins were detected through the addition of 20 μL of an Amplex UltraRed mixture (prepared according the manufacturer’s instructions (ThermoFisher)). Turnover of the substrate was monitored at 27°C on a Synergy^TM^ HT Microplate Reader. All values were determined in triplicates.

For the competition ELISA, mTFP1 was immobilized on plates through its SBP-tag as described above and incubated for 1 h with 20 μl of 100 nM FLAG-tagged anti-mTFP1 DARPins alone or in combination with 500 nM HA-tagged competitor DARPins (non-binding DARPin E3_5 as well as either of the two anti-mTFP1 DARPins). For detection of the bound FLAG-tagged DARPins, wells were incubated for 1 h each with a primary mouse anti-FLAG antibody (Sigma-Aldrich #F3165, 1:5,000) and secondary goat anti-mouse-AP antibody (Sigma-Aldrich # A3562, 1:10,000), followed by the addition of 20 μl per well of a 3 mM p-nitrophenyl phosphate solution (Sigma-Aldrich, #71768). Absorption was measured 30 min after incubation at 37°C at 405 nm. All measurements represent technical triplicates.

The FA assay was performed as described previously (Brauchle et al., 2014) using black non-binding 96-well plates (Greiner). Constant amounts of mTFP1 (15 nM) were titrated with a dilution series of DARPins (four replicates) and the fluorescence anisotropy was measured on a Tecan M1000 equipped with a suitable anisotropy module. The K_D_ was determined by fitting the data with a non-linear fit using GraphPad Prism.

### Crystallization, data collection and structure determination

All crystallization experiments were carried out with 15 mg/ml protein in sitting-drop vapor diffusion experiments. mTFP1/DARPin 1238_E11 crystals in space group C2 grew after 2 days at 4°C in 10% PEG4000, 20% glycerol and 0.02 M of L-glutamate, glycine, DL-alanine, L-lysine, DL-serine. mTFP1/DARPin 1238_E11 crystals in space group P6_5_22 grew within one week in 0.1 M imidazole pH 7.0 and 30% 2-methyl-2,4-pentanediol at 20°C. Plate-like crystals of DARPin 1238_G01 in space group I4 appeared after one week in 2 M NaCl, 10% PEG10,000 at 20°C and grew to their final size within two weeks. Crystals of isolated DARPin 1238_E11 in space group P2_1_ grew after 2 months in 0.2 M (NH_4_)_2_SO_4_, 0.1 M MES pH 6.5, 20% PEG8000 at room temperature. Rod-like DARPin 1238_G01 crystals in space group P2_1_2_1_2_1_ grew within 2 days in 0.2 M NaF, 20% PEG3350 at 20°C. mTFP1/DARPin 1238_E11 crystals were directly frozen in liquid nitrogen. DARPin 1238_E11 and DARPin 1238_G01 crystals were cryo-preserved by addition of ethylene glycol to a final concentration of 20 % (v/v) and flash-cooled in liquid nitrogen. All measurements were done at the SLS beamlines X06DA and X06SA (Swiss Light Source, Paul Scherrer Institute, Switzerland) at 100 K. All data were integrated, indexed and scaled using the XDS software package (Kabsch, 2010a; Kabsch, 2010b). Data collection statistics are summarized in Supplementary Table S1. The structures were solved by molecular replacement using the crystal structure of anti-IL4 DARPin (PDB ID: 4YDY) and mTFP1 (PDB ID: 2HQK) (Ai et al., 2006) as search models with the program Phaser (McCoy et al., 2007). Model building, structure refinement and model validation were performed with Coot (Emsley and Cowtan, 2004), PHENIX (Adams et al., 2002), Refmac5 (Murshudov et al., 2011) and Molprobity (Chen et al., 2010), respectively. Refinement statistics are summarized in Supplementary Table 1. The atomic coordinates have been deposited in the RCSB Protein Data Bank and are available under the accession code 6FP7, 6FP8, 6FP9, 6FPA and 6FPB, respectively.

### Plasmid construction

All the eukaryotic expression plasmids were generated by specific PCR amplification and standard restriction cloning. Briefly, anti-mTFP1 DARPins, including the N-terminal His-tag and the C-terminal Flag-tag, were PCR amplified from the bacterial expression constructs and inserted into pmCherry (Hybrigenics, France). For the DARPins-YPet fusions, the mCherry coding sequence was replaced by the YPet coding sequence by standard PCR and restriction cloning. The additional polyisoprenylation CAAX peptide (GGGRSKLNPPDESGPGCMSCKCVLS) of the human K-Ras oncogene protein, or the whole human HISTH2BJ (histone H2B) coding sequence were inserted at the C-terminus of the DARPins-YPet fusion constructs for generating the membrane-(DARPins-YPet-CAAX) or nuclear (DARPins-YPet-H2B) tethering DARPins. The mitochondrial bait mito-mTFP1, containing an N-terminal anchor sequence from the human CISD1 protein (the first 59 amino acids) fused to the N-terminus of mTFP1, was generated from pcDNA4TO-mito-mCherry-10xGCN4_v4 (Addgene plasmid 60914 (Tanenbaum et al., 2014)) by substituting the mCherry coding sequence with that of mTFP1 and substituting the 10xGCN4_v4 tags with 1xGCN4_v4 tag. mTFP1-CAAX and HIST2BJ-mTFP1 were cloned into CMV expression vectors (pmKate2-N, Evrogen,) replacing mKate2 with mTFP1 coding sequences. For the mTFP1-Rab5c fusion construct, both coding sequences of mTFP1 and zebrafish (*Danio rerio*) Rab5c were cloned in this order into pcDNA3 (Invitrogen) by standard PCR and restriction cloning. All constructs were verified by sequencing. Plasmid maps and oligonucleotide sequences for PCR and cloning are available upon request. A schematic representation of the fusion constructs is provided in Supplementary Figure S4 and their resulting fusion protein amino acid sequences are given in Supplementary file S1.

### Cell cultures, transfections and imaging

HeLa S3α cells were maintained in Dulbecco’s modified Eagle’s medium supplemented with 10% foetal calf serum, 100 IU penicillin and 100 μg streptomycin per ml. One day before transfection, cells were seeded on glass cover slip placed into a 24 well plate at a density of 50,000-100,000 cells/well.

Transfections were carried out with 1 μg of total DNA (500 ng for each construct or with empty expression plasmid) and 3 μl of FuGENE ® HD Transfection Reagent (Promega), according to the manufacturer’s instructions. 24 hours post transfection, cells were fixed in 4% paraformaldehyde, stained with Hoechst 33342 (Invitrogen) and mounted on standard microscope slides with VECTASHIELD® (Vector Laboratories Inc. Burlingame, CA).

Confocal images were acquired with a Leica point scanning confocal “SP5-II-MATRIX” microscope (Imaging Core Facility, Biozentrum, University of Basel) with a 63x HCX PLAN APO lambda blue objective and 1-2x zoom.

## Acknowledgments

We would like to thank Simon Ittig, Christoph Kasper, Marilise Amstutz of T3 Pharmaceuticals for their generosity to host one of us (MAV). We also thank the Imaging core facility of the Biozentrum for their assistance, all the members of Affolter lab for helpful discussions and Bernadette Bruno, Gina Evora and Karin Mauro for their great help in the media kitchen. We further acknowledge all current and former members of the High-Throughput Binder Selection facility at the Department of Biochemistry of the University of Zurich for their contribution to the establishment of the semi-automated ribosome display that resulted in the generation of the anti-mTFP1 DARPin binders, especially Thomas Reinberg, Valerio Berardi and Jonas Kapp. We also acknowledge the beamline staff at Swiss Light Source (Villigen, Switzerland) for their excellent support.

The work in the Affolter lab was supported in parts by grants from SNF and SystemsX (MorphogenetiX); the work on the DARPin selection was funded by SNF (grant 310030B_166676 to AP) and the University of Zurich.

---

**Supplementary Figure S1**

**Structural analysis of the mTFP1/DARPin complex.**

**(A)** Sequences of selected DARPins. The sequences of DARPin 1238_E11 and DARPin 1238_G01 are shown. The top row indicates the consensus sequence used in the library with randomized positions indicated as X (randomization to all amino acids but Cys, Pro and Gly) and Z (randomization to Asn, His or Tyr), highlighted in black frames. In the lower lines, only residues that differ from the consensus sequence are printed. **(B)** Superposition of the DARPin 1238_G01 (grey) and DARPin 1238_E11 (rainbow-colored from N-(blue) to C-terminus (red) (C_α_ r.m.s.d. 0.3 Å). (C) Superposition of the DARPin 1238_E11 (grey) and DARPin 1238_E11 in complex with mTFP1 (lime) (C_α_ r.m.s.d. 0.45 Å), minor differences are only observed for the two N-Cap helices. **(D)** Superposition of the mTFP1/DARPin 1238_E11 and the GFP/ DARPin 3G61 complex (PDB ID: 5MAD) (Hansen et al., 2017). mTFP1 (teal), DARPin 1238_E11 (grey), GFP (green) and anti-GFP DARPin 3G61 (rainbow coloured, from N-terminus blue to C-terminus red) are shown in cartoon representation. The mTFP1 chromophore is shown as a sphere. **(E)** Superposition of the mTFP/DARPin 1238_E11 and the GFP/DARPin 3G124nc complex (PDB ID: 5MA8) (Hansen et al., 2017). mTFP1 (teal), DARPin 1238_E11 (grey), GFP (green) and anti-GFP DARPin 3G124nc (rainbow coloured, from N-terminus blue to C-terminus red) are shown in cartoon representation. The mTFP1 chromophore is shown as a sphere.

**Supplementary Figure S2**

**Intracellular binding of anti-mTFP1-DARPins**

Confocal images of HeLa cells transiently transfected with pH2B-mTFP1 alone (first row) or with the combination of either pH2B-mTFP1 and pCMV-DARPin 1238_E11-mCherry (second row) or pH2B-mTFP1 and pCMV-DARPin 1238_G01-mCherry (third row), respectively; pmTFP1-CAAX alone (fourth row) or the combination of either pmTFP1-CAAX and pCMV-DARPin 1238_E11-mCherry (fifth row) and or pmTFP1-CAAX and pCMV-DARPin 1238_G01-mCherry (sixth row). The first column represents the mTFP1 bait (green), the second column is the mCherry DARPin channel (red), the third column is the overlay of the first two channels, showing the nuclear colocalization of the H2B-mTFP1 bait (second and third rows) or membrane colocalization of pmTFP1_CAAX bait (fifth and sixth rows) with the respective DARPins. The fourth column represents the nuclear Hoechst staining (blue) and the fifth column is the merge of all three channels (with the scale bar in white (15 μm) on the bottom right corner). Images were taken 24 hours post transfection. Transfected constructs are indicated at the right of each row and the different channels are indicated at the top of each column. The figures are from a representative experiment, performed at least three times.

**Supplementary Figure S3**

**Intracellular binding of anti-mTFP1-DARPins-YPet**

Confocal images of HeLa cells transiently transfected with pCMV-DARPin 1238_E11_YPet alone (first row) or in combination with pH2B-mTFP1 (second row) or mito-mTFP1 (third row). The same transfection scheme is presented for the DARPin 1238_G01-YPet alone (fourth row) or in combination with H2B-mTFP1 (fifth row) or mito-mTFP1 (sixth row). The first column represents the mTFP1 channel (green), the second column is the YPet channel (magenta), the third column is the overlay of the two channels, showing the colocalization of the DARPins-YPet fusion to the different mTFP1 baits, the fourth column represents the nuclear Hoechst staining (blue) and the fifth column is the merge of all three channels (with the scale bar in white (15 μm) on the bottom right corner). Images were taken 24 hours post transfection. Transfected constructs are indicated at the right of each row and the different channels are indicated at the top of each column. The figures are from a representative experiment, performed at least three times.

**Supplementary Figure S4**

**Schematic representation of the fusion constructs**

The transcriptional elements (enhancer, promoter and poly (A) adenylation) of the different mammalian expression vectors are depicted as grey filled boxes. The different protein coding modules are represented as coloured block arrows, while the resulting fusion protein is depicted as a solid black arrow inside the modules. Full maps and sequences are available upon request.

**Supplementary file S1**

Amino acid sequences of the fusion proteins used in the manuscript. The different modules are coloured as depicted in Supplementary Figure S4

**Supplementary Table S1**
Statistics on diffraction data and refinement of DARPin 1238_E11, DARPin 1238_G01, and mTFP1/DARPin 1238_E11 complexes.

